# In a temporally segmented experience hippocampal neurons represent temporally drifting context but not discrete segments

**DOI:** 10.1101/338962

**Authors:** J. H. Bladon, D. J. Sheehan, C. S. De Freitas, M. W. Howard

**Affiliations:** Center for Memory and Brain, Boston University, Commonwealth Ave., Boston, MA, 02215; Graduate Program for Neuroscience, Boston University, Commonwealth Ave., Boston, MA, 02215

## Abstract

There is widespread agreement that episodic memory is organized into a timeline of past experiences. Recent work suggests that the hippocampus may parse the flow of experience into discrete episodes separated by event boundaries. A complementary body of work suggests that context changes gradually as experience unfolds. We recorded from hippocampal neurons as male long evans rats performed 6 blocks of an object discrimination task in sets of 15 trials. Each block was separated by removal from the testing chamber for a delay to enable segmentation. The reward contingency reversed from one block to the next to incentivize segmentation. We expected animals to hold two distinct, recurring representations of context to match the two distinct rule contingencies. Instead, we found that overtrained rats began each block neither above nor below chance but by guessing randomly. While many units had clear firing fields selective to the conjunction of objects in places, a significant population also reflected a continuously drifting code both within block and across blocks. Despite clear boundaries between blocks, we saw no neural evidence for event segmentation in this experiment. Rather, the hippocampal ensemble drifted continuously across time. This continuous drift in the neural representation was consistent with the lack of segmentation observed in behavior.

**Significance Statement:** The neuroscience literature yet to reach consensus as to how hippocampal firing fields support the organizing of events across time in episodic memory. Initial reports of hippocampal activity focused on discrete episodes within which representations were stable, and across which representations remapped. However, it remains unclear whether this segmentation of representations is merely an artifact of cue responsivity. More recently, research has shown that a proportion of the population codes for temporal aspects of context by exhibiting varying degrees of drift in their firing fields. Drift is hypothesized to represent a continually evolving temporal context, however it is unclear whether this drift is continuous or is also a mere artifact of changing experiences. We recorded from the dorsal hippocampus of rats performing an object discrimination task that involved contexts that were segmented in time. Overtrained rats were unable to anticipate the identity of the upcoming context, but may have used context boundaries to their advantage. Event segmentation theory predicts that hippocampal ensembles would alternate between behaviorally-relevant segments. Contrary to these predictions, animals showed weak evidence of context segmentation, even across blocks with different reward contingencies. Hippocampal ensembles showed neither evidence of alternating between stable contexts nor sensitivity to context boundaries, but did show robust temporal drift.

## Introduction

Episodic memory refers to the vivid recollection of a specific event situated in a unique place and time (E Tulving & Madigan, 1970). In his description of episodic memory, Endel Tulving emphasized that a crucial distinction between episodic and semantic memory is that episodic memories are temporally dated, or are remembered in relation to other events across time. The hippocampus is essential for episodic memory and is thought to mediate this function by binding events to a representation of spatial and temporal context (H. Eichenbaum, Yonelinas, & Ranganath, 2007; Endel Tulving, 1972). Indeed the hippocampus is specifically involved in both remembering the temporal order of past events and making associations across a temporal gap in both humans and animals (Dede, Frascino, Wixted, & Squire, 2016; Fortin, Agster, & Eichenbaum, 2002; Kesner, Hunsaker, & Gilbert, 2005; Scoville & Milner, 1957). Hippocampal ensembles are theorized to represent a ‘cognitive map,’(O’Keefe & Nadel, 1978) that acts as a contextual or relational scaffold onto which events may be bound together for later retrieval (Davachi, 2006).

There are two complementary models for how spatiotemporal context is structured in the hippocampus. Event segmentation theory suggests experience is segmented across time into discrete situational contexts (Figure 1 “Event Segmentation”) (Baldassano et al., 2017; DuBrow, Rouhani, Niv, & Norman, 2017; Muller & Kubie, 1987; Zacks, Tversky, & Iyer, 2001). Temporal context theory suggests that the hippocampal representation of context evolves continually (Figure 1 “Temporal Context”) (Howard & Eichenbaum, 2013; Howard, Fotedar, Datey, & Hasselmo, 2005). There is behavioral, neuroimaging and animal neurophysiological evidence consistent with both theories.

**Figure 1.**
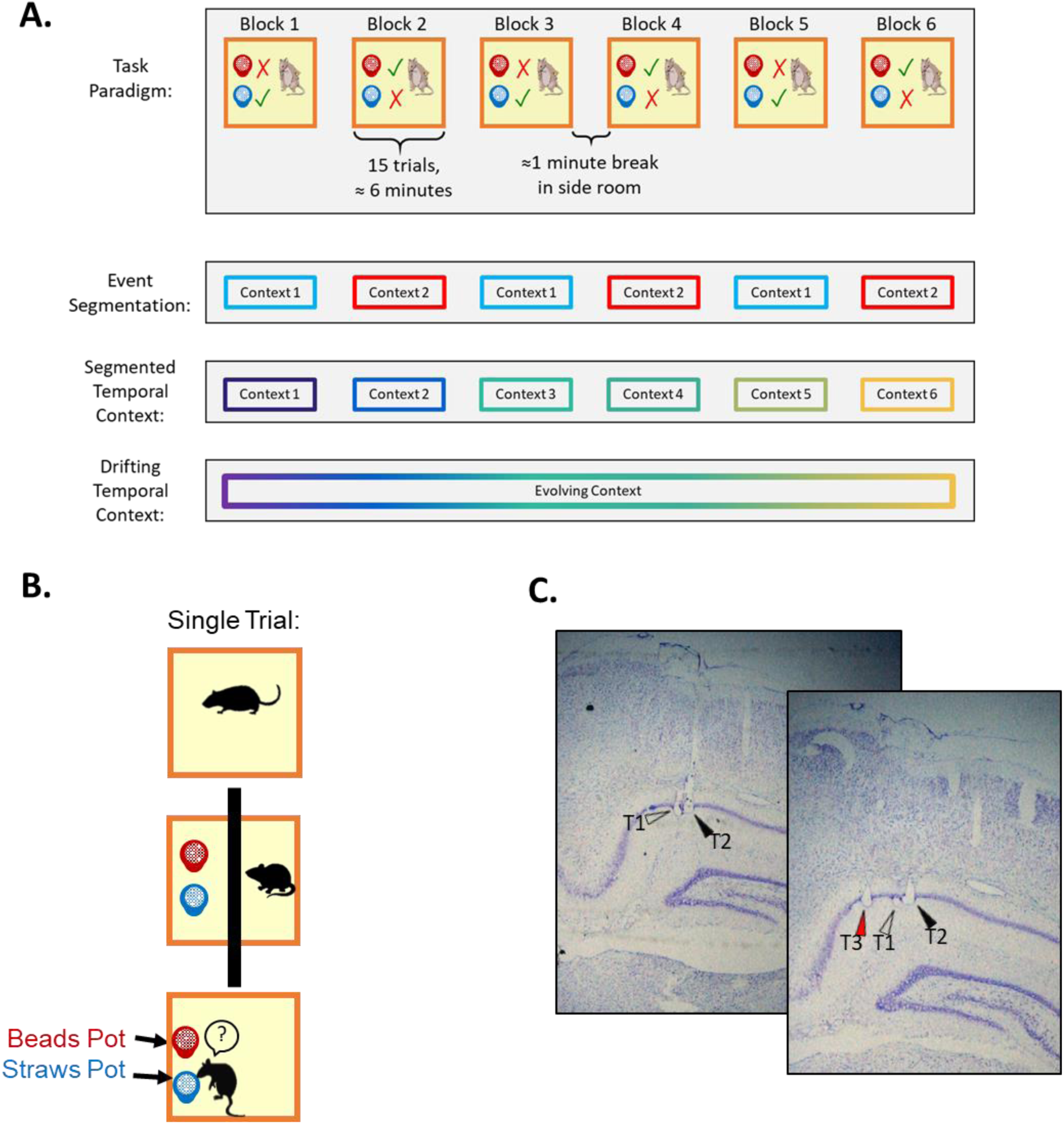
Task Design. **A**. Rats performed a blocked object discrimination task in which the reward contingency was held constant for 15 trials in a row and then reversed for the next 15. Each block took roughly 6 minutes, and each delay between blocks was fixed at 1 minute. **B**. Each trial consisted of three phases, a short inter trial interval, the pot setup phase, and the sample/choice phase. **C**. Recording locations of three individual tetrodes. Arrowheads indicate final tetrode locations in the pyramidal layer of dorsal CA1.

Consistent with event segmentation theory, behavioral and neuroimaging evidence suggests that experiences are segmented in time. Studies on human recall have found that memories are organized by discrete situational contexts (Baldassano et al., 2017; DuBrow et al., 2017; Muller & Kubie, 1987; Zacks et al., 2001). Abrupt changes in environmental or contextual cues across time can cause a behavioral separation in memory traces (Ezzyat & Davachi, 2011; Sols, Dubrow, Davachi, & Fuentemilla, 2017; Zacks et al., 2001). Hippocampal BOLD activity in humans, and ensemble activity in rodents increases when a border between contexts is perceived, as if the hippocampus parcellates experience into contextual chunks (Baldassano et al., 2017; Bulkin, Sinclair, Law, & Smith, 2018; DuBrow et al., 2017; Mack, Love, & Preston, 2016; Place, Farovik, Brockmann, & Eichenbaum, 2016).

Although event segmentation has yet to be explicitly studied in rodents, electrophyisological evidence consistent with event segmentation can be found in the hippocampal representation of space (but see Bulkin et al. 2018 for a recent exception). In agreement with event segmentation theory, hippocampal place cells generate separate maps across different spatial environments (Brandon, Koenig, Leutgeb, & Leutgeb, 2014; Komorowski, Manns, & Eichenbaum, 2009; Leutgeb, Leutgeb, Moser, & Moser, 2005; Wills, 2005). In addition, hippocampal ensembles generate separate maps across a variety of explicit contextual designations that occur in the same physical space (Kobayashi, Nishijo, Fukuda, Bures, & Ono, 1997; Markus, Barnes, McNaughton, Gladden, & Skaggs, 1994; Smith & Bulkin, 2014; Wills, 2005). In a particularly clear example, neural segmentation across two physical contexts was found to develop as rats learned to discriminate between the two contexts (Komorowski et al., 2013a, 2009). These studies suggest that hippocampal ensembles segment continuous experience to support discrimination between different spatial and behavioral contexts.

Consistent with temporal context theory, behavioral and neuroimaging experiments suggest a continuous temporal signal that supports the organization of memories in time. The temporal contiguity effect describes the tendency for subjects to bind together unrelated events that occurred together in time, and has been shown across timescales in both neural and behavioral datasets (Folkerts, Rutishauser, & Howard, 2018; Howard, Youker, & Venkatadass, 2008; Kahana, 1996; Manning, Polyn, Baltuch, Litt, & Kahana, 2011; Zaromb et al., 2006). Conversely, a continually evolving hippocampal signal also allows for segregation of related events that occur far apart in time (Cai et al., 2016; Manns, Howard, & Eichenbaum, 2007). Human fMRI data show the representational similarity of hippocampal BOLD signals during recall of events reflects a broad continuum of relatedness that maps onto the temporal and spatial proximity of those events (Deuker, Bellmund, Navarro Schröder, & Doeller, 2016; Hsieh, Gruber, Jenkins, & Ranganath, 2014; Jenkins & Ranganath, 2016; Nielson, Smith, Sreekumar, Dennis, & Sederberg, 2015; Schapiro, Kustner, & Turk-Browne, 2012; Schapiro, Turk-Browne, Norman, & Botvinick, 2016).

There is a large body of neurophysiological evidence both in rodents and in humans that supports the temporal context theory. The classic view of hippocampal place cells suggests the hippocampal map represents the same spatial context across long periods of time, extending for weeks to months (Thompson & Best, 1990). However these initial studies focused on only a small population of cells specifically sought out for their stability. Numerous studies have reported slow changes in the representation of place across extended time (Folkerts et al., 2018; E. A. Mankin et al., 2012; Emily A. Mankin, Diehl, Sparks, Leutgeb, & Leutgeb, 2015; Manns et al., 2007; Mau et al., 2018; Paz et al., 2010; Rubin, Geva, Sheintuch, & Ziv, 2015; Ziv et al., 2013). Crucially, these reports consistently show that there are both stable and drifting units in the active ensemble, suggesting hippocampal ensembles are tracking both the similarities and differences in experiences across time. Indeed, multiple reports have found that while a minority of units are active across many days, there remains a stable population from which one could decode the animal’s spatial context and location within that context across large temporal lags (Mau et al., 2018; Rubin et al., 2015; Ziv et al., 2013). These reports span temporal scales and show both stability and drift on the order of seconds, minutes, hours, and days. Moreover, these phenomena have been observed across a wide variety of recording methodologies suggesting this is a feature of the system, rather than a measurement artifact (Cai et al., 2016; E. A. Mankin et al., 2012; Manns et al., 2007; Mau et al., 2018).

There is behavioral, neuroimaging and animal neurophysiology evidence consistent with both temporal drift and event segmentation, and the two hypotheses are not mutually exclusive (Figure 1 “Segmented Temporal Context’). However, these two bodies of work have never been directly compared in the same preparation. The goal of this experiment was to examine whether or how these two forms of context representation are observed in a task that leverages a temporally defined context. This task paradigm involves two distinct behavioral contexts that are separated by a behavioral boundary. The hippocampus is hypothesized to map these contexts into discrete event representations as proposed by event segmentation theory, and to also contain a slowly drifting signal as proposed by temporal context theory.

## Experimental Methods

In order to determine how the hippocampal map segments similar experiences that occur across minutes, rats performed a task in which distinct behaviors were reinforced in different temporal blocks of trials in the same spatial environment. The boundary between blocks was cued by shuttling the animal to a separate chamber for 1 minute, but there were no overt cues to signal the behavioral context at the time of the choice behavior. A representation of temporal context that changes continuously across blocks would be behaviorally suboptimal in this experiment (Figure 1 “Drifting Temporal Context”). Rather, the strategy to maximize reward would be to segment the experiment into behaviorally meaningful contexts by using the boundary cue (Figure 1 “Event Segmentation”).

During performance of this task, we recorded extracellularly from dorsal CA1 ensembles. As in earlier work that showed evidence for spatial event segmentation (Komorowski et al., 2013, 2009), rats were presented with pairs of pots containing unique odors and digging media and the rat was rewarded for choosing the correct pot from the pair. The identity of the rewarded pot was consistent for each block of 15 trials after which the 1-minute boundary cue was imposed; for the next 15 trials the other pot was rewarded. The event segmentation hypothesis predicts two stable mappings of objects and places, one for each rule condition (Figure 1 “Event Segmentation” panel). In contrast, the temporal context hypothesis predicts a continuous decorrelation of neural representations that was unaffected by the blocked structure of the experience (Figure 1 “Drifting Temporal Context” panel). A representation validating both theories might involve a new but stable context instated at the start of each block of trials (Figure 1 “Segmented Temporal Context” panel).

### Subjects

Subjects were 5 male Long-Evans rats (Charles River) weighing between 350 and 450 grams and between the ages of 6 months to 1 year for the duration of the experiment. All animals were single housed and maintained on a 12 h light/dark cycle (lights on 8:00 A.M. to P.M.). Behavioral training and testing were conducted exclusively during the light phase. Animals were maintained at a minimum (85%) of their *ad libitum* feeding body weight during all behavioral training and testing periods. Procedures were conducted in accordance with the requirements set by the National Institutes of Health and Boston University Institutional Animal Care and Use Committee (IACUC).

#### Behavioral Apparatus

The behavioral training and testing environment was a custom-built wood apparatus (40 l × 60 w × 40 h cm) consisting of a 40 cm × 40 cm box, and a 20 cm × 20 cm side alleyway. The objects consisted of identical circular terra cotta pots (10 cm high with an internal diameter of 9 cm), each with their own unique digging media and odors (e.g., purple beads with grapefruit scent). The pots were distinguishable only by their scent and digging media, requiring the animal to overtly sample before choosing to dig. In order to prevent the animals from being guided by odor of the Froot Loop (Kellogg’s) cereal reward, finely crushed Froot Loops were sprinkled into all digging media.

### Surgery

Anesthesia was induced by inhalation of 5% isoflurane (Webster Veterinary Supply) in oxygen and then a stable plane was maintained at 1.5%-3% throughout the entirety of surgery. Before surgery animals were injected with the analgesic Buprenex (buprenorphine hydrochloride, 0.03 mg/kg i.m.; Reckitt Benckiser Healthcare), and the antibiotic cefazolin (330 mg/ml i.m.; West-Ward Pharmaceutical). The skin of the animal’s head covering the skull was shaved and cleaned with alcohol swabs before then being placed in a stereotaxic frame (Kopf). A longitudinal incision was made to expose the skull and the bone, and underlying fascia was cleared in order to gain access to stereotaxic coordinates and locations for anchoring screws. Animals were implanted with microdrives containing 18–24 independently drivable tetrodes targeting the dorsal pole of the CA1 cell layer of the hippocampus (centered at 3.6 mm posterior and 2.6 mm lateral from bregma). Finally a screw was placed above the cerebellum to serve as a ground signal. Each tetrode was composed of four 12 um RO 800 wires (Sandvik Kanthal HP Reid Precision Fine Tetrode Wire; Sandvik). Tetrodes were plated with non-cyanide gold solution, via electrolysis in order to reduce impedance to between 180 and 220 k**Ω**. At the conclusion of the surgery, all tetrodes were gradually lowered ~0.5 − ~1.5 mm into tissue. Upon recovery from anesthesia, animals underwent post-operative care for 3 days and received doses of Buprenex and cefazolin, as described above, two times a day (12 hour intervals). Animals were allowed to recover 1 week before behavioral testing commenced.

### Neural Recordings

Electrophysiological recordings for this project were collected on a 96 channel OmniPlex D Neural Acquisition System (Plexon). Each channel was amplified on head-mounted preamps and then amplified again for total of 1000x to 10,000x before being digitized at 40 kHz. Spike data were bandpass filtered from 200 Hz to 8.8 kHz and local field potentials from 1.5 Hz to 400 Hz. Spike channels were referenced to a local electrode in the same region in order to remove both movement-related and ambient electrical noise. That local reference electrode was then referenced to ground and provided the LFP signal in the region. Action potentials of neurons were detected via threshold crossing and then sorted later using Offline Sorter (Plexon). Cineplex Studio (Plexon) was used for capturing behavioral tracking data, and Cineplex Editor (Plexon) was employed to enter event markers and to verify animal tracking data. Between recorded training sessions tetrodes were advanced at a minimum of 40 μm and positioned based on visual inspection of spike clusters in order to maximize neural unit yield. Tetrodes were allowed to settle after turning over a period of days to prevent contamination of neural signals with tetrode drift.

### Histology

Upon completion of behavioral testing, rats were anesthetized with <5% isoflurane in oxygen. Anatomical recording sites were confirmed by creating a small lesion in the brain tissue by passing a 40 μA current until the connection was severed (generally 2–8 seconds). Immediately after completion of electrolytic lesions, animals received an overdose injection (interperitoneal) of Euthasol (Virbac AH) and upon cessation of breathing were immediately transcardially perfused with ice cold 0.5% potassium phosphate buffered saline followed by 5% phosphate buffered formalin (VWR). Brains were then removed and placed in additional 5% formalin phosphate for at least 36 hours. Brains were then submerged in 30% sucrose for cryoprotection until sectioning into 40 μM thick sections via cryostat (CM 3050s; Leica Biosystems). Brain sections were processed using standard Nissl staining protocol in order to visually confirm tetrode-recording sites (Figure 1c).

### Animal Training and Task

Once each animal recovered from surgery they were initially trained to dig for Froot Loop (Kellogg) bits in an aloe-scented pot over a cloves scented pot (both in sand). Once they reliably dug in the aloe pot and refrained from digging in the cloves pot, training began in the blocked-reversal task. Each training and recording session consisted of 6 blocks of 15 trials for a total of 90 trials. Once the rat reached a criterion of 70% correct across a given session, recording commenced. Some sessions were terminated early (the shortest session was 85 trials) due to lack of motivation by the subject. Within each 15 trial block the reward contingency was set so that one pot always had food and the other pot did not. Each trial started with the insertion of a divider so that the experimenter could place the pots in a pseudorandom position. Once pots were placed, the divider was removed and the animal was allowed to sample each pot, but only allowed to dig in one. Upon digging, the unchosen pot was immediately removed and the animal was allowed to dig until he found reward or gave up because he chose incorrectly. After either completion of reward consumption or a 3 second delay following pot removal, the rat was shuttled to the far half of the chamber and the divider replaced. The next trial commenced immediately until trial 15 was reached. After the last trial of each block, the rat was shuttled into the side alley to wait for 60 seconds. After the break, the reward contingency was reversed and the next block of 15 trials was performed.

#### Quantitative & Statistical Analyses

All analysis of the collected data were performed using custom scripts from MATLAB (MathWorks). ANOVAs were performed using the ‘anovan’ function in MATLAB under a standard type 2 sum of squares. For individual unit analyses, peri-event histograms were generated from 120 to 160 millisecond bins and smoothed using a moving average of a three bin span. All trials were included in peri-event rasters including those in which the rat responded incorrectly, but only the first sample in each trial is shown. Right and left samples correspond to each of the two pseudorandomized locations of the reward pots. Error bars on firing rates were calculated using the standard error of the mean. Spatial firing rate plots were generated using a 3 cm pixel size, and then convolved with a Gaussian smoothing kernel with a standard deviation of one pixel. Selectivity was evaluated by calculating a selectivity score below, where n represents the set of trial-types(two in the case of object and position, four in the case of object by position), and *λ* represents the mean firing rate for that event type. *λpref* represents the trail type with the largest firing rate.

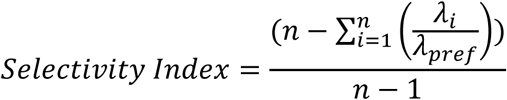

Significance was determined by generating a null distribution of selectivity scores after randomizing the trial identity of each sample. Only units that passed a 99% significance threshold from 10,000 boots were considered to code a particular dimension (Keene et al., 2016).

For waveform and spike rate drift metrics, average waveform amplitude and spike rate was estimated for each 10 second bin spanning the whole recording session. We then measured the average absolute Euclidean distance and absolute difference in firing rate between each bin to all other bins. From that matrix, we regressed the values as a function of distance from the diagonal but excluding the diagonal to obtain an average waveform or spike rate distance as a function of temporal lag between bins for each cell. Units with an average firing rate of >10 Hz across the whole recording session were assumed to be interneurons and were removed from all population analyses.

For population analyses, trial rate vectors were constructed for each cell by averaging the firing rate across the two seconds surrounding the first sampling event on each trial, and then z-normalizing the rate across trials for each individual neuron. All trials were included in these analyses, so as to reveal any performance effects as well as for statistical reasons. A z-transform was employed to prevent overreliance on highly active units or under reliance on sparsely active units. A population vector correlation matrix was generated for each rat by calculating the Spearman correlation of the population vector for each trial to each other (Figure 6B). That correlation matrix was then averaged across all sessions to generate the ‘super rat’ matrix observed in Figure 6A. All measures of drift were constructed by first generating a mean value for each session onto which statistics were performed, and the mean +/− SEM across sessions was plotted. Bootstrap permutation tests were performed as described with a standard 10,000 randomized samples in which the group index was randomized without replacement. Bayes factors were obtained by inputting summary statistics into an online engine accessible at http://pcl.missouri.edu/(Liang, Paulo, Molina, Clyde, & Berger, 2008).

## Results

### Rats performed a Temporally Blocked Object Discrimination Task

We recorded from 768 cells across 25 sessions from 5 rats each implanted with a 24-tetrode hyperdrive aimed at dorsal CA1 (Figure 1C). After removing putative interneurons and keeping only units with an average firing rate of below 10 Hz and with at least one spike for at least 5 sampling events we obtained 642 putative pyramidal cells. Rats were trained to perform a blocked object discrimination task designed to segment memory into six 15 trial blocks (Figure 1A). Within each block of 15 trials, one of the two distinguishable pots contained hidden reward. After each 15 trial block, a temporal delay signaled the end of a block; the reward contingency was reversed to the other pot for the subsequent block of trials (See Figure 1 “Event Segmentation”). Between blocks the rat was shuttled into a side chamber for 60 seconds so that the end of a block was signaled not only by an increased trial duration *per se* but also by an intervening experience. Individual trials took on average a little under 30 seconds to perform (26.0 +/− 0.40 sec), each block of trials took about 6 minutes (6.04 +/− 0.56 min), and a session lasted roughly 45 minutes (45.44 +/− 1.1 min). Each session contained an unequal but roughly similar number of sampling events at each item and position, however some animals showed a slight bias towards sampling the item on one side of the maze more often. Furthermore, we restricted our analyses to the first sampling event on each trial, as the identity of the object on the first sample remained pseudorandomized per task design. Only after the sample had terminated and a response was offered did the behavior systematically change towards rejecting the incorrect object and digging in the correct pot.

### Rats performed as though each block of trials was a new episode

Following pre-training on simple pot discrimination, rats took roughly a week to reach a criterion of 70% correct within a given session. Recording began after criterion was reached. Once trained, all rats performed similarly to each other (Mean across rats=80% +/− 1.97%, ANOVA, F(4,20)=0.1), and all rats performed similarly across days (Mean across days=79% +/− 0.52% ANOVA, F(7,17)=1.61) and blocks within each day (Mean across blocks=79%+/−1.13% ANOVA, F(5,144)=1.73). Typically, errors were concentrated around the beginning of each block, but were not restricted to the beginning of the recording session (Figure 2). We generated two alternative hypotheses regarding behavior. We hypothesized that the rats would recognize the block transitions as a cue to change their response strategy, and would begin each block by switching their response towards digging in the now-correct pot. Alternatively, rats could have ignored the boundary cue. In this case the animals would begin each block by erroneously perseverating with the (now-incorrect) response from the previous block. We evaluated these hypotheses at the group level, but also at the individual animal level (if some rats respected the contextual cues while others did not, the group could perform at chance on average despite none of the individual rats performing at chance). To investigate both hypotheses, we compared each rat’s performance for each trial across blocks to chance by performing a binomial test on each animal. A two-sided binomial test asks whether the probability of correct responses was *either* above *or* below chance. Contrary to both hypotheses, only one rat performed different from chance for either of the first two trials in each block before a bonferonni correction (Figure 2, Two Sided Binomial Test, Rat 1 Trial 1: 30% uncorrected binom p=0.03, trial 2: 38%, binom p=0.24: Other rats Trial 1: 45.6%+/− 5.7%, minimum p=0.09, trial 2: 50.4%+/− 6.21%, minimum p= 0.31). By trial 5 all rats were performing significantly above chance (Figure 2, trial 5: 81.5%+/− 4.38%, All Binom. P<0.05, trials 6 to 15: 90.7% +/− 1.2%, All Binom. p< 0.005). Rats appeared to begin each block by impulsively digging in the first pot they encountered, as the probability of rejecting the first encountered pot on the first trial of each block was lower than for the last 8 trials of each block (Trial 1 reject rate mean=0.32 +/− 0.038%, trials 11:15 reject rate mean=0.42 +/− 0.014%, ranksum p<0.05). Thus, rats appeared to begin each block of trials by guessing, and then rapidly learning the new rule contingency. The performance of each rat on each block could be described as either a recency-weighted averaging over recent experiences. Alternatively, they may have perceived the block delay as a new context but the behavioral contingency of that context was unknown to the rat. Both hypotheses account for the chance performance following the inter-block delay and also the learning of the new reward contingency within block (Figure 2 Right). However, an interpretation consistent with recency weighted averaging predicts a neural representation consistent with drifting temporal context (Figure 1), whereas recognition of the boundary but not anticipation of the rule predicts a neural representation consistent with either a segmented temporal context, or event segmentation (Figure 1).

**Figure 2.**
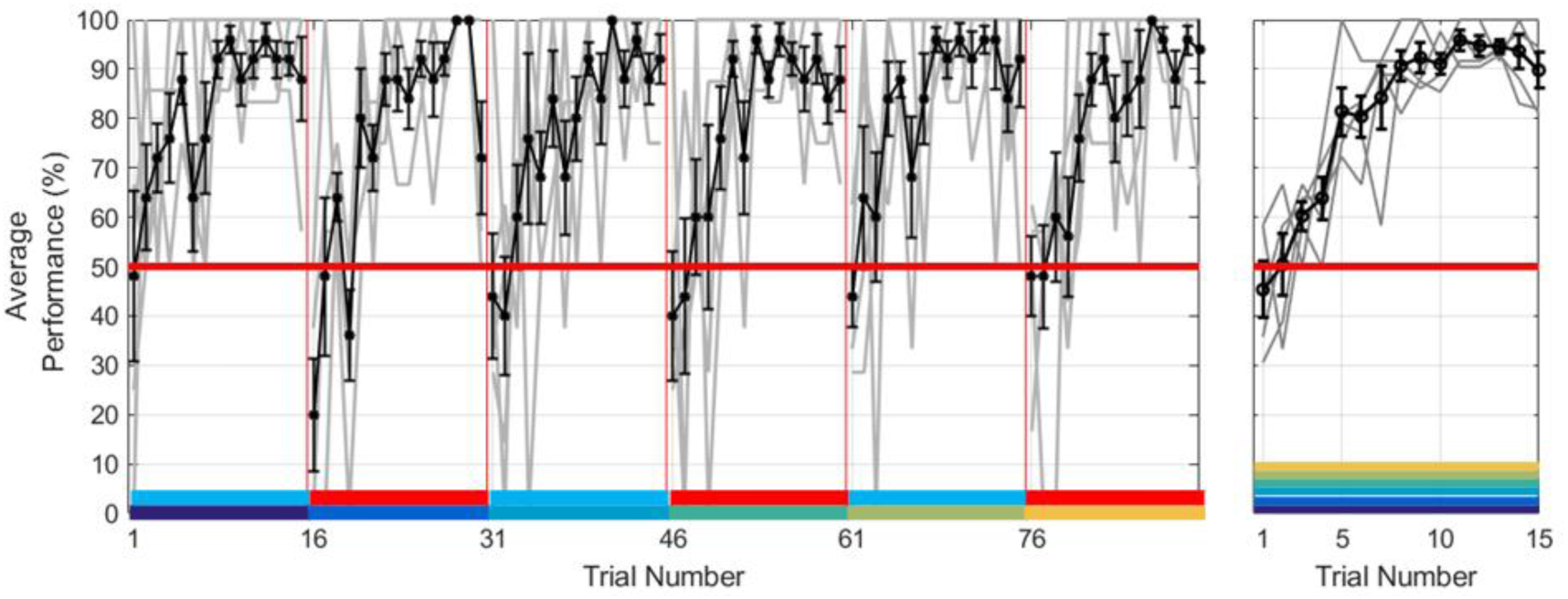
Rat’s behavior did not anticipate the reversal in reward contingency across blocks. **Left**: Mean +/− SEM performance throughout each recording session. Gray lines represent individual rats. Rats began each block in a session at chance regardless of the position of that block in the session. (Colored Bars represent changing context per Figure 1) **Right**: When all blocks were concatenated (colored blocks stacked), no rat performed better or worse than chance at the beginning of each block. Gray lines represent individual rats while black represents mean across all rats. Red line is chance performance. All rats behaved as though each block represented a novel context in which to learn the rule contingency (colored bars represent evidence towards ‘temporal context’ as denoted in Figure 1).

### Single units replicated prior findings of object and position selective fields, but were impacted by context

Single unit activity was observed by generating spatial heat plots (Figure 3), and peri-event rastergrams, and histograms centered on pot-sampling events (Figure 4). Consistent with previous reports, a large proportion of place cells had firing fields where the objects were presented (Figure 3, cells 1, 3, 5, and 6). When event-locked firing was examined, there was a large overlapping population of units whose firing fields discriminated between sampling events. Some units had firing fields consistently discriminated item regardless of position (Figure 4 top row, 43 cells (8% of putative pyramidal cells), showed an object selectivity score greater than 99% of 10,000 bootstrap permutations). There were also units whose fields were specific to one position (Figure 3, 106 cells (17% of putative pyramidal cells) showed a position selectivity score greater than 99% of 10,000 bootstrap permutations) as well as those specific to one item in one place (Figure 4 bottom row, 80 cells (12% of all putative pyramidal cells) showed object by position selectivity greater than 99% of 10,000 bootstrap permutations). Upon further investigation firing fields showed clear dependence on the shifting context (McKenzie et al., 2014) (Figure 5).

**Figure 3.**
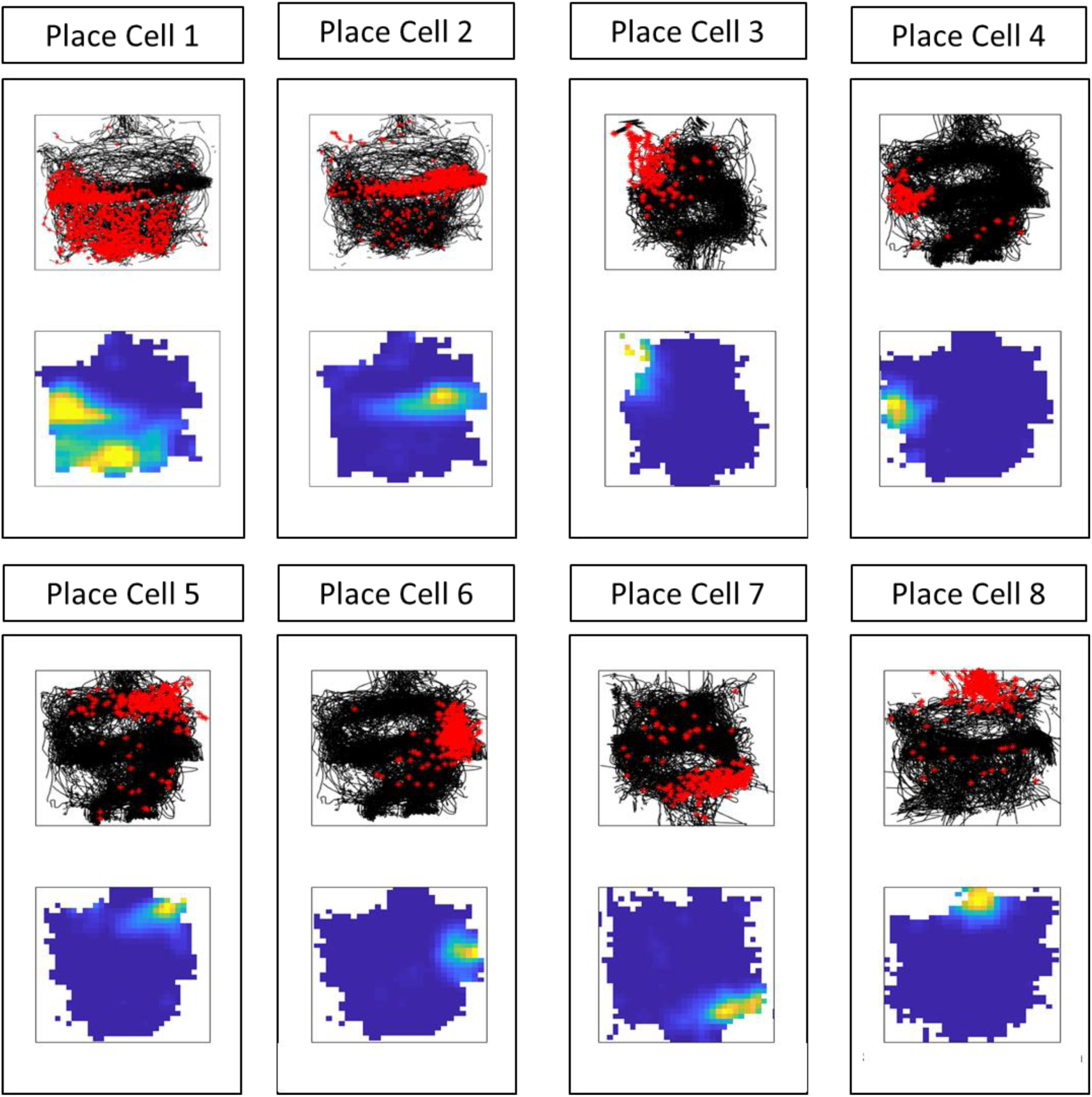
Many units showed spatially localized firing fields. **Top plots**: Black lines denote the running trajectories of the rat, while red dots denote the locations of spikes recorded from one unit. **Bottom plots**: spatial heat maps of binned-firing rates for each unit. Yellow denotes maximal firing, blue denotes minimal firing.

**Figure 4.**
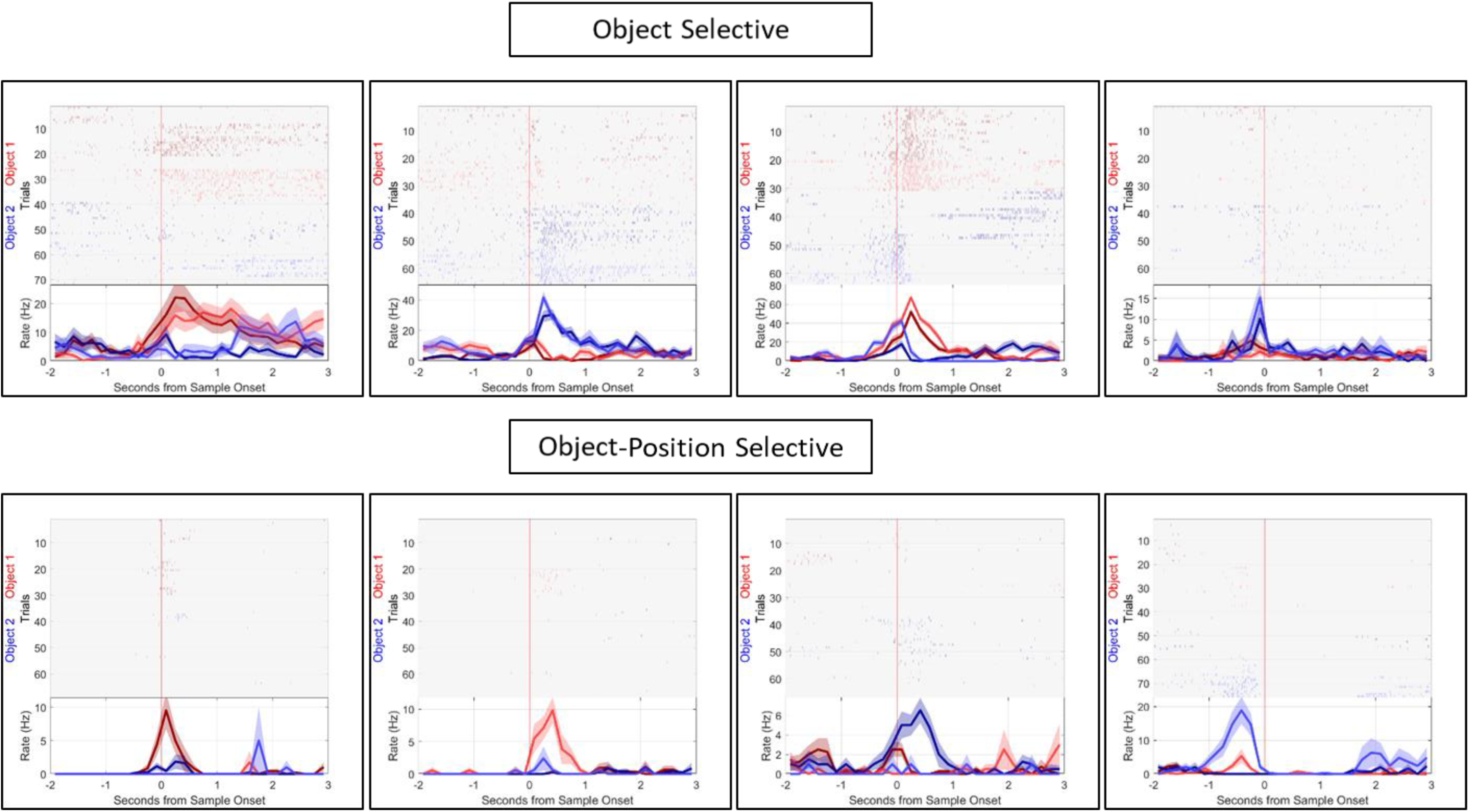
Many units showed object-specific firing, (top four) and conjunctive object -position firing (bottom four). Peri-event rasters and histogram plots centered on object sampling. Time refers to seconds from sample onset, and rate refers to firing rate in Hz. We found some units to be object selective (red vs. blue) regardless of the position of the object (light shades v dark shades), as well as some objects to be selective to one object-position combination.

**Figure 5.**
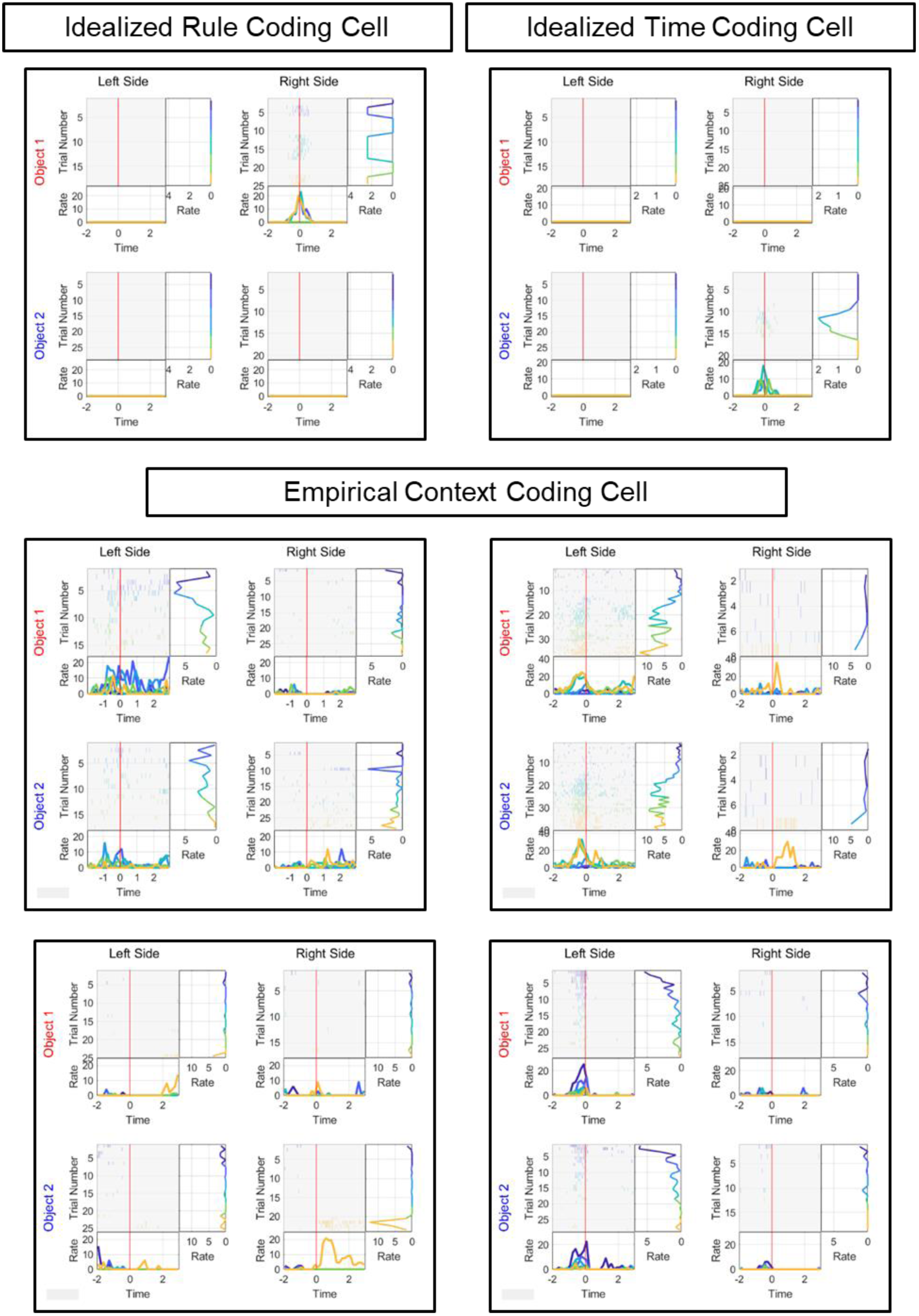
Many cells had context dependent firing fields. **Top**: Idealized firing fields of dCA1 units in the temporally blocked object discrimination task. Each color in rasters and line plots represent samples in one block of trials (See Figure 1A and Figure 2). **Top Left:** Ideal cell that responds selectively during one object-position combination, and does so only during blocks 2, 4, and 6 when object 2 is rewarded. **Top Right**: Idealized object-position conjunctive cell that responds maximally during block 4. Each idealized cell provides information about the object, position, and temporal structure of the task. **Bottom:** Empirical cells showed firing fields that seemed to be centered on a small contiguous block of trials, suggesting temporal drift. No empirical cells showed firing fields that alternated with blocks, showing a lack of evidence for event segmentation.

### A drifting contextual representation replicates previous findings and is uncorrelated to waveform drift

Many units showed firing fields that were modulated by the changing context. The temporal context model and the event segmentation model each make a strong prediction for how context may modulate hippocampal firing fields (Figure 5). While putative pyramidal cells maintained the same spatial and object selectivity across blocks of trials, their rates showed obvious changes across contexts that seemed to resemble predictions from the temporal context model (Figure 5, bottom). Figure 5 contains plots that include all trials including error trials, and each trial was color coded based on block (Figure 5, Color Scheme: Figure 1 “Segmented Temporal Context”). Briefly, event segmentation theory predicts that in this experiment firing fields should alternate across blocks so that units’ response should be similar in the same the behavioral context. Conversely, temporal context theory predicts that in this experiment firing fields should change continuously without regard to the alternating behavioral contexts. For example, the first example cell (Figure 5) had a firing field that was selective to leftward samples of object 1, and was most robust for the first two blocks of trials. That is, this unit fired across two blocks of trials with different behavioral contexts and then ceased firing despite the repetition of those behavioral contexts.

### Gradual Changes in firing rate were uncorrelated with electrode stability

To exclude the possibility that cells were drifting due to acquisition error, spike clusters from each tetrode were carefully examined as a function of time. First, tetrodes that included waveform clusters that obviously appeared not stationary with regard to time we excluded (Manns et al., 2007). Then, drift in spike amplitude was correlated to drift in spike rate in the remaining population of cells. For each cell, the Euclidean (4 dimensional) distance was measured between the average spike amplitudes of each 10 second bin in the session and then the distances were regressed against their bin lag to obtain a measure of waveform drift. The same was performed for absolute difference in average firing rate at the same 10-second bins to obtain a measure of firing rate drift. While there was a distribution of spike amplitude drift rates and activity drift rates, there was no relationship between the two measures (Figure 6, Spearman’s Rho, r^2^ (642) = 0.0606, p>.05).

**Figure 6.**
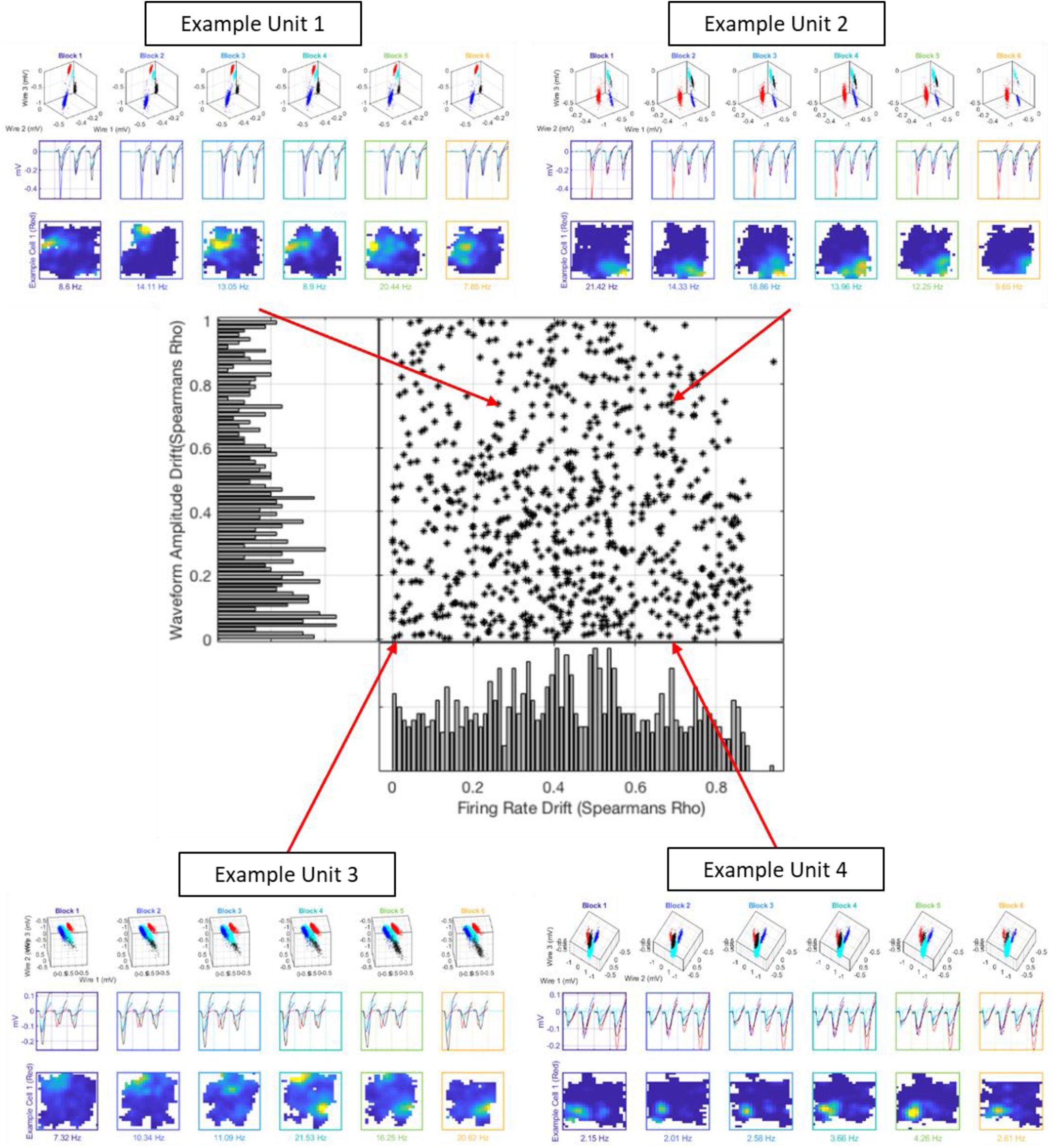
Waveform drift is unrelated to spike rate drift. For each cell we measured the average spike rate and waveform shape for each 10 second bin across the recording session, and then measured the similarity within each metric at bins of increasing lag (see methods). Cells showed a wide distribution of spike-rate drifts (vertical line plot at bottom), and a wide distribution of spike shape drift rates (horizontal bars at left). However, the spike-rate drift was unrelated to spike shape drift. Each example represents one cell at each corner of the scatter plot. Each unit cluster and waveforms are represented in red, and other units on that tetrode are included. Peak firing rates are listed below place plots for each unit during each block. Firing rate and waveform drift rates for each example unit, respectively: Unit 1: 0.26, 0.73; Unit 2: 0.69, 0.73; Unit 3: 0. 07, 0.01; Unit 4: 0.70, 0.08.

These data revealed a wide spectrum of firing rate drift rates, as well as a wide spectrum of spike amplitude drift rates. However, as there was no relationship between these two distributions, the single unit data suggested that individual units exhibit a continuous spectrum of drift rates across time that is not due to recording artifact.

### Ensemble activity suggests hippocampal patterns slowly drift

To supplement the analysis on individual units, population analyses were used to determine whether hippocampal ensembles showed evidence for event segmentation, a drifting temporal context, or both. Event segmentation predicts that the hippocampus generates two stable representations, one for each rule condition (Fig 7A top left). This would manifest in alternating high correlations between blocks of the same rule condition, and low correlations between blocks of opposing rule condition. A drifting temporal context predicts that ensembles slowly change, and new representations would be continually generated (Figure 7A top right). This would manifest as a slow fall in the correlation between blocks at increasing temporal lag. We calculated the Spearman correlation of the population vector from each trial to all others to generate a correlation matrix for each rat (Figure 7B, Methods). We then averaged that matrix across all rats and sessions to generate a grand mean correlation matrix (Figure 7A Bottom). The empirical pattern of ensemble activity showed strong support for drift, characteristic of temporal context (Figure 7A right) but no apparent evidence that representations from past blocks with the same reward contingency were repeated (Figure 7A left). Correlation values were greater at the beginning and the end of the session, but this may have been due to the lack of an acclimatization period at the start, and the fact that some sessions were shorter than others (Monaco, Rao, Roth, & Knierim, 2014). Importantly, the correlation values did not fall smoothly at increasing distances from the diagonal of the matrix. This was likely due to behavioral variables, such as where the rat was during the sampling event, and which pot the rat was sampling. Correlation matrices were reorganized to control for these variables. First trials were sorted by position such that the first and fourth quadrants contained trial pairs in the same position. This revealed a strong ensemble code for space, as exemplified in rats 2 and 4 (Figure 7B) where higher correlations were clustered at same position comparisons. The events were then sorted by object and lastly by time, to reveal correlations that fell more smoothly with increased distance from the diagonal (Figure 7B right). Furthermore, temporal proximity was evident across object and place representations in some rats as evidenced by yellow streaks parallel to the diagonal of the matrix, but away from the diagonal of the matrix (Figure 7B right, rats 1 and 4). These stripes signify that temporally proximal trials that differed in object or place were represented more similarly than those that were farther apart in time. Thus, there was also a portion of the ensemble that tracked trial lag, but not object or position.

**Figure 7.**
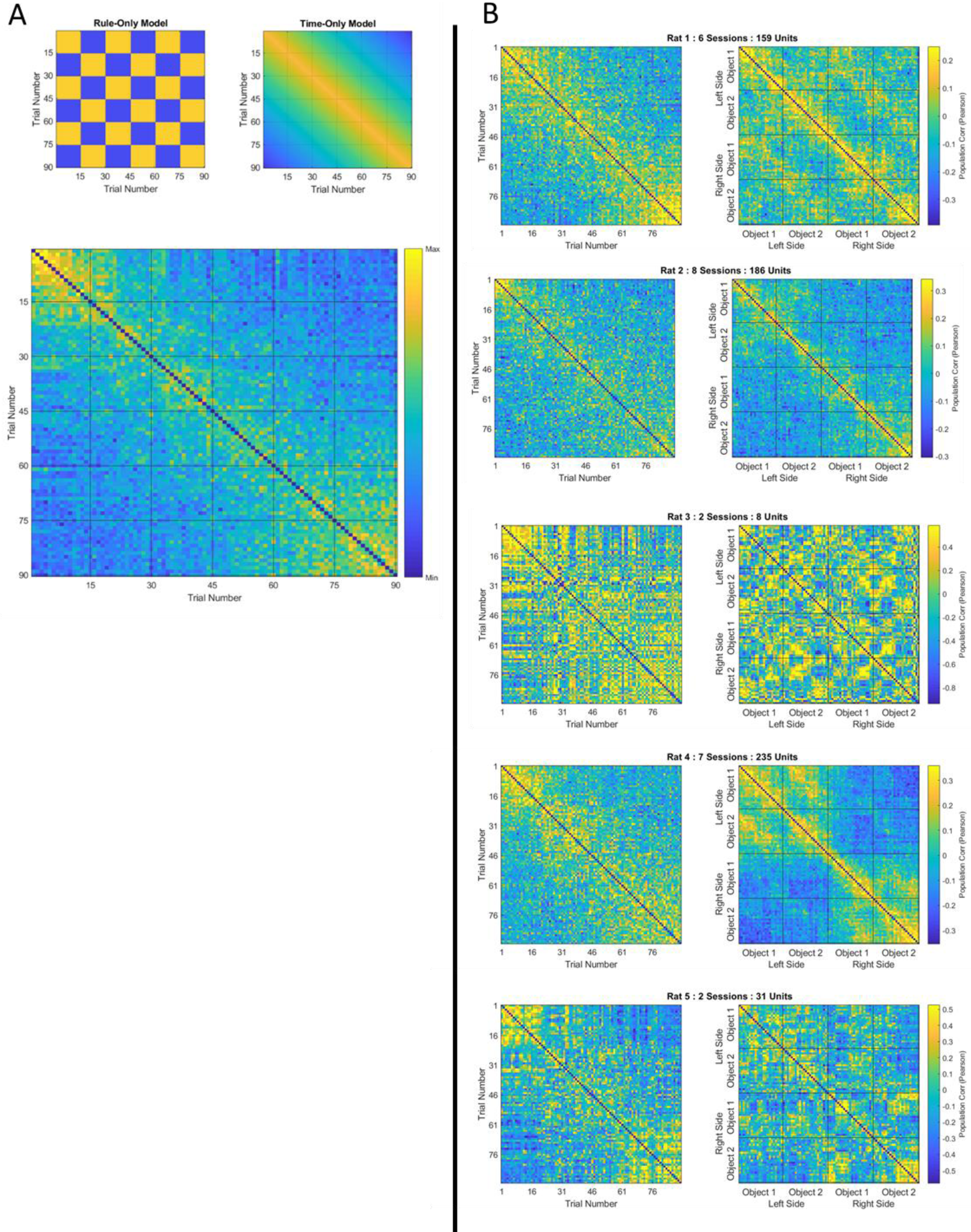
Drift, and not a representation of the repeating rule conditions was apparent in correlation matrices. **Figure7A**. Observed correlation matrix suggests continuous temporal drift. **Top left**: Predicted correlation matrix if ensembles represent a context code consistent with event segmentation. **Top right**: Predicted correlation matrix if ensemble code is consistent with drifting temporal context. **Bottom**: Empirical correlation matrix averaged across animals more closely resembles predicted matrix under the temporal context hypothesis. **Figure 7B**. Trial-by-Trial correlation matrices for each rat also show temporal drift. **Left**: Matrices were sorted by trial number as was done in Figure 7A. **Right**: Matrices sorted by position, then by object, then by trial number. Position coding can be seen as higher correlations in the top left and bottom right quadrants of each matrix. Object coding can be seen as nested quadrants within each position quadrant. Note that once trials were sorted by object and place that high correlations still clustered adjacent to the eye of the matrix, indicating that ensembles were more similar between trials at closer temporal proximity. Similarity was calculated by generating z-normalized firing rate vectors for each cell across sampling events, and then calculating the spearman’s rho between the population vectors on each trial. The color scale is equivalent across the two matrices for each rat, but a different scale was used for each rat.

### Hippocampal Populations showed drift across blocks

The effects observed in the correlation matrices were then quantified. If hippocampal ensembles reflected the two rule states, hippocampal activity would be more similar between blocks that shared a rule condition (at lag 2, and 4), versus those that involved opposing rule conditions (at lag 1, 3, and 5) (Figure 8 top left). Conversely, if hippocampal ensembles tracked time, then hippocampal activity would become progressively less similar between blocks at increasing lag (Figure 8 bottom left). The overall population correlation between blocks consistently fell as the block lag grew (Figure 8 Right, purple line) (across all trials slope=−0.025, Observed slope exceeded all 10000 perms, permutation mean −2.38×10^−7^, σ=0.0034). The representation for the same behavior (e.g. same object, position, and response) also progressively decorrelated with block (Figure 8, Right, yellow line) (Permutation test: slope=−0.048, the observed slope exceeded all 10,000 bootstrapped permutations, permutation x=−1.5×10^−5^, σ=0.0073). Correlations for the same item, position, and response remained higher than those across all sample types at each block lag (Bonferonni corrected Ranksum test: Lag 1: Within object, position, response mean correlation = 0.16, across mean correlation =0.03, p<1×10^−4^, Lag 2 within mean =0.08, across mean =−0.02, p<0.001, Lag 3 within mean =0.03, across mean =−0.06, p<0.001, Lag 4 within mean =0.01, across mean =−0.07, p<0.001, Lag 5 within mean =−0.05, across mean =−0.08, p<0.05) suggesting a robust code for objects and places even though both populations drifted. Finally, when these results were reevaluated after only using units with the most stable waveform clusters (most stable quartile, see Figure 4) the population still significantly decorrelated with increasing temporal lag, and at a similar rate (Slope=−0.038, Observed slope exceeded all 10,000 perms, permutation mean 1.12×10^−4^, σ=0.0087). Finally, the representation of each delay in between blocks also progressively decorrelated with increasing lag (Figure 8 Black line, Slope of correlation vs. lag=−0.076, observed slope exceeded all 10,000 perms, perm mean=−3.6 × 10^−3^, σ=0.017). Therefore, all empirical curves replicate previous findings of a temporal context coded in conjunction with a stable code for items, places, and behavior. Note that deviations from the smoothly decreasing curves are small-any contribution from discrete event coding would have to be much smaller than the contribution due to gradually-changing temporal context. Moreover, event segmentation would predict that the deviations from a smooth curve should be consistent from one type of comparison to the other, resulting in parallel curves (Figure 8, top left). However, to the extent there were deviations in the different empirical curves, they were not systematic across the type of comparison. Thus, consistent with behavior, there was no evidence that hippocampal populations represented blocks in discrete segments based on the two rule conditions.

**Figure 8.**
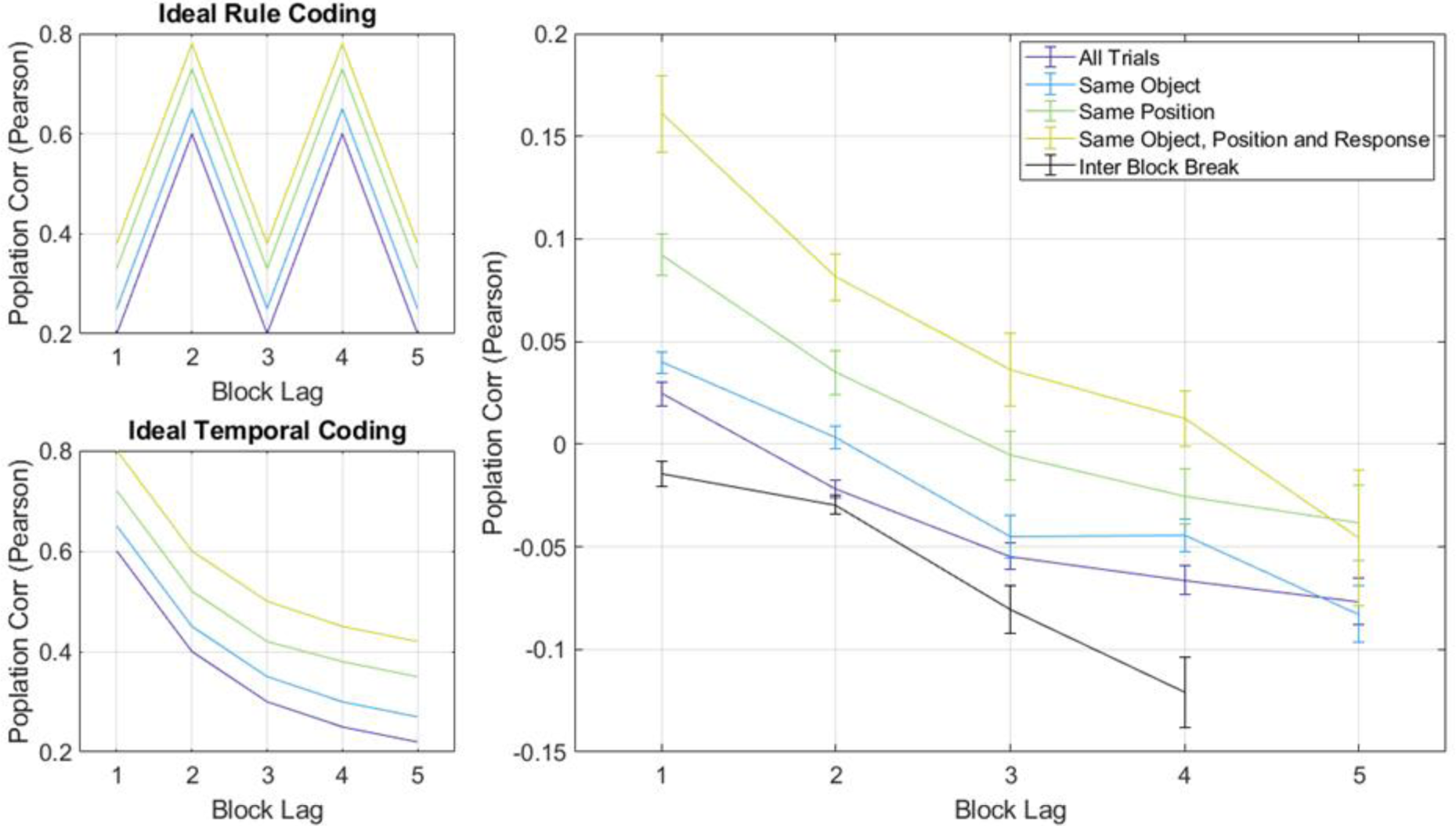
Ensembles reflected context code that continually changed across blocks and did not recur. **Top left**. Ideal curves under event segmentation hypothesis. Event segmentation predicts high correlations between blocks of the same rule condition at lags 2 and 4. **Bottom Left**. Ideal curves under the contextual drift hypothesis. Contextual drift predicts a monotonic decrease in each curve. **Right**. Observed curves for ensemble correlations fell as the block lag between the trials grew. This was true for the overall population (purple line) as well as in the ensemblecoding the same object (blue) or position (green) or same object, position and behavioral response (yellow), and finally the intervals between the blocks (black line). All curves show a systematic decrease from small to large lags with no obvious alternation. To the extent that there are fluctuations in the curves, these fluctuations were small and not consistent across different types of comparisons.

Overall hippocampal ensembles showed strong temporal drift with no evidence that past blocks of the same rule condition used the same code. This was true for the overall population, but was also found in conjunction with a code for places, objects, and object-place conjunctions. Furthermore the code for objects and positions remained across all blocks. Thus even though hippocampal populations drifted from block to block, there remained a code for objects and positions that persisted throughout the recording. Temporal drift also persisted during the inter-block-delays, where there was also no evidence for an alternating neural structure.

### Hippocampal Populations showed drift within block

The foregoing analyses demonstrate that the hippocampal ensemble changed gradually across blocks on the scale of tens of minutes. This subsection examines changes in hippocampal representation within a block on the scale of about a minute. To replicate previous findings, population correlations were compared between individual trials (Fig. 9)(Manns et al., 2007). There was significant temporal drift in the overall representations across trials (Figure 9 purple line, observed slope=−6.8×10^−3^, observed data exceeded all 10,000 perms, perm. x =−8.16 ×10^−6^, σ=8.4×10^−4^). Temporal drift within a block was apparent after controlling for object, (Figure 9 blue line, Observed slope= −8.5×10^−3^, observed data exceeded all 10,000 perms, perm. x=6.98 ×10^−6^, σ=1.1×10^−3^), position (Figure 9 green line, Observed slope= −7.1×10^−3^, observed data exceeded all 10,000 perms, perm. x =−1.58 ×10^−5^, σ=9.0×10^−4^), and object, position, and response (Figure 9 yellow line, Observed slope= −0.013, observed data exceeded all 10,000 perms, perm. x =−4.65 ×10^−7^, σ=1.3×10^−3^). Drift was also apparent across exclusively correct trials (Not shown, Observed slope= −0.0095, observed data exceeded all 10,000 perms, perm. =−1.32 ×10^−5^, σ=1.7×10^−3^) suggesting that this is not an artifact of the learning curve observed in Figure 2. This drift occurred individually in the overall population of each rat tested except for Rat 3 (Figure 8B, Rat 3 included 2 sessions with 12 units total, observed slope=-0.011, observed data exceeded 716/1000 perms, perm x=−1.24 ×10^−4^, σ=0.0076). This failure to observe robust drift in Rat 3 is perhaps because Rat 3 only contributed a total of 12 units over 2 sessions.

**Figure 9.**
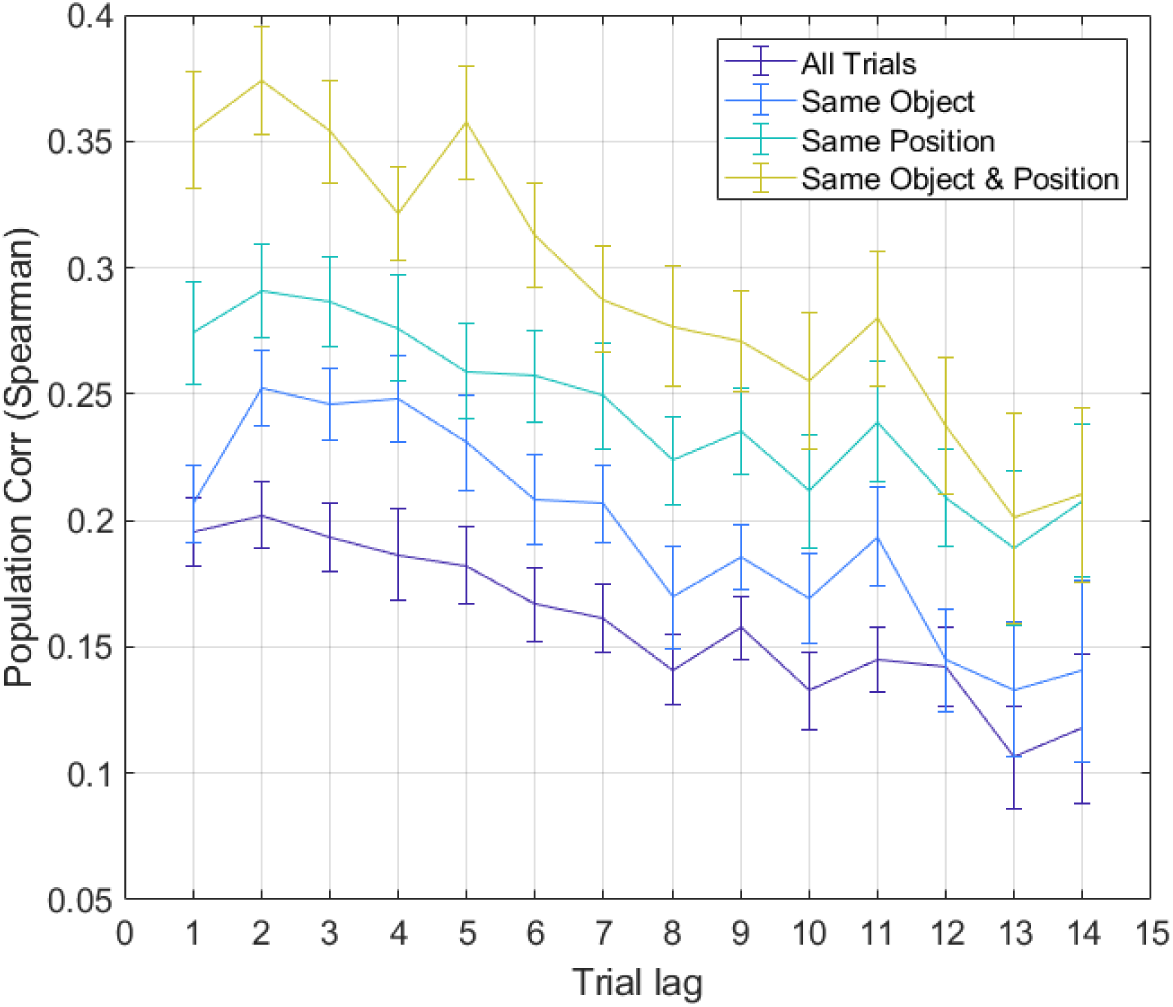
Ensemble correlations within a block reliably fell as trial lag increased. This effect was evident both in the overall population activity (purple line), as well as when we controlled for object (blue), position (green), or object, position, and response (yellow). The overall correlations across lags were higher after controlling for object, position, and object-position conjunctions, revealing a population code for objects and positions.

These data replicate previous findings that show drift is observable across individual trials. Drift on the order of seconds was observed in the overall code, as well as the code for objects, positions, and object-position conjunctions. Thus, reliable population drift was observed even across iterations of the same behavior and context, and over a scale of seconds.

### Hippocampal shifts between blocks can be accounted for by time

We observed robust drift in the hippocampal population occurring both across blocks of trials as well as on a trial to trial basis. However, it is possible that in conjunction with drift within each block, there exist shifts in population state at the transitions between blocks (see Figure 1, Segmented Temporal Context vs. Drifting Temporal Context). This would manifest as an increase in the population drift rate across blocks over what was observed within a block. To isolate this possible effect, trial pairs within a block (Figure 10 left, blue line) were examined separately from those that spanned a block transition (Figure 10 left, red line) after controlling for object and position effects. We observed significant drift both within each block as well as across blocks (Within block observed slope=−.395, observed data exceeded all 10,000 perms, perm mean=−7.66×10^−4^, σ=0.055. Across block observed slope=−0.279, observed data exceeded all 10,000 perms, perm =−5.276 ×10^−4^, σ=0.0543). After controlling for trial lag the block transitions induced a separation in representations of contexts as other studies have suggested (Baldassano et al., 2017). Because quantification of segmentation is inversely related to that of drift in this instance, choosing segmentation as the alternative hypothesis leaves drift as the null. Standard statistical testing was therefore problematic, as it provides only positive evidence for one alternative hypotheses over the null. Therefore, we used a Bayes Factor to directly compare the likelihoods of segmentation and drift and find positive evidence for the more likely hypothesis, equally assessing the alternative and null. The correlation between trials in the same block was significantly greater than that in adjacent blocks after controlling for trial lag (JZS Bayes T-Test yielded a Bayes Factor strongly in favor of a difference in means, odds ratio 400:1). This suggests that the transition between blocks induced a separation between representations of trials that happened in different blocks consistent with event segmentation.

**Figure 10.**
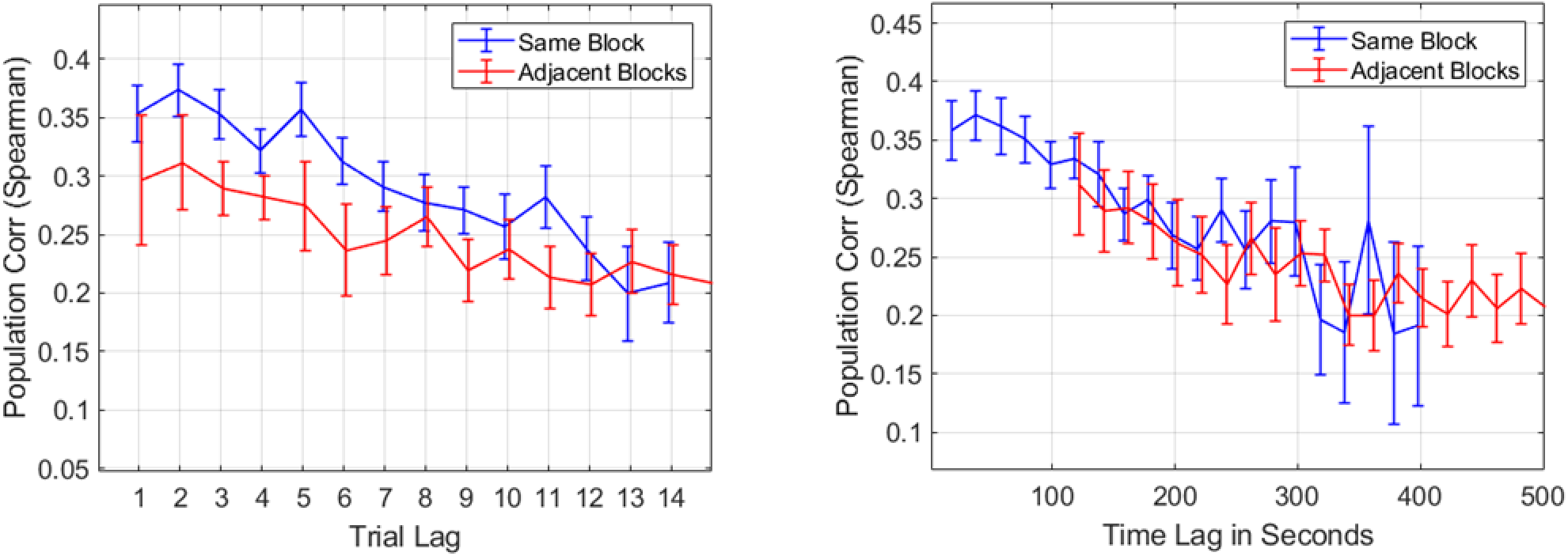
Time fully accounts for shifts in population state across blocks. **Left**. Population vectors within a block (blue) were more similar than those in adjacent blocks (red) when trial lag was considered. Alone, this might have been evidence for event segmentation. **Right**. Population vectors within block (blue) were *not* more similar than those in adjacent blocks (red) when time was considered. Population vector correlations were lower in adjacent blocks, but time was sufficient to account for this decrease. Thus, there was no evidence for event segmentation between blocks above and beyond the change attributable to the temporal delay between blocks. Population vectors were constructed in the same manner as Figure 10.

This could either be because the boundary cue induced a separation in hippocampal representations, or that the separation in representations was merely a consequence of the extended time between the two trials spanning the delay. To address whether the extended temporal lag fully accounted for the apparent boundary effect on the hippocampus, we plotted the correlation between trials by their *temporal* lag and then organized pairs by whether there was a block transition between them (Figure 10 right). When we compared trial pairs in the same block with those in adjacent blocks in this way, we found that the block break caused no more reduction in the population vector correlation than would be expected by elapsed time (JZS Bayes T-Test yielded a Bayes factor strongly in favor of a single mean, odds ratio 31:1). Thus the separation of trials spanning a block transition observed in the left figure was completely eliminated by accounting for elapsed time. This was also true after removing trials at the beginning of the block when performance was poor (Trials 4:15 of each block only, JZS Bayes T-Test yielded a Bayes Factor strongly in favor of a single mean, odds ratio 15.9:1) removing the possibility that the representation of context only shifted after the animal had switched behavioral strategy. Thus, even though every animal reliably alternated their choices between blocks and no rat perseverated into the next block, there was no evidence that hippocampal code segmented experience any more than what would be expected from elapsed time.

## Discussion

In this experiment we sought to determine how the hippocampus codes for behaviorally-relevant context in a task in which animals were required to segment experience in time to distinguish between two rule contingencies. Event segmentation predicts that animals would have parcellated experience into discrete episodes based on contextual boundaries; temporal context predicts that the hippocampal representation should provide a continuous relational metric that progressed consistently across time. We provided rats with a temporally blocked task such that an event boundary (removal from the testing chamber) cued a reversal in the reward contingency. Importantly, the context boundary was a transient cue that signaled a change in which object was rewarded, but was not an explicit cue that would be available to the animals as they made the choice.

Interestingly, well-trained animals neither perseverated across cued block transitions, nor did they anticipate the rule reversal. Instead, every rat tested began every block at chance, frequently digging in the first pot he encountered. Thus, while no rat learned that there were two alternating rule conditions, they all benefitted from the boundary cue by changing their behavior across the block boundaries. Hippocampal units showed selectivity for the position, object, and object-position conjunction of sampling events while also showing sensitivity to the changing context. Indeed, the ensemble code showed strong evidence for temporal drift both across blocks and within each block. Conversely, there was no evidence that the ensemble code was more similar for blocks that shared a rule contingency. Furthermore, there was no greater separation in the hippocampal representation than was expected by the passage of time across the event boundaries that marked block transitions.

### The methodological design could have prevented a segmented neural representation

The behavioral results suggest that while no rat learned to anticipate the reversal of the reward contingency between blocks that nonetheless they all were sensitive to the boundaries between blocks. There are multiple interpretations of these curious results. One clue was that the probability of rejecting the first pot encountered was significantly lower at the start of each block, suggesting the rats began a block with no a-priori preference. Perhaps rats forgot the past rule condition during the block transition and entered the box naïve, only remembering that one of the pots contained reward, but not which pot. Alternatively, they may have merely grown impatient during the delay, and therefore were unwilling to reject either pot. Previous data suggest this behavior may reflect a hippocampal dependent cognitive flexibility, rather than mere impatience. Numerous studies on blocked alternative choice behaviors revealed that hippocampal lesions caused rats to perseverate more after switches in rule contingency or reward location (Hsiao & Isaacson, 1971; Kimble & Kimble, 1965). Importantly, each of these studies involved behavior that was fixed for a large block of trials followed by an abrupt change in reward contingency. Considering our animals showed no consistent perseveration across blocks, these results suggest that the dorsal hippocampus might have been involved in the small savings the rats exhibited at the beginning of each block. Indeed, the drift we observed in the hippocampal representation aligns well with the observed behavior as if the dorsal hippocampus played a role in supporting the flexible changes in responses across time.

Further support for this interpretation may be found in studies on the recency effect observed in human and animals that also show a moderately hippocampal dependent strengthening of memory for recent list items (Kesner, Crutcher, & Beers, 1988). Under this interpretation, the long delay caused a weakening of memory that contributed to the rats’ uncertainty at the beginning of each block. This uncertainty might have manifested as a reduced rejection rate. These two accounts suggest that even though animals were unable to track the two recurring rule conditions, their behavior in this task may have been supported by hippocampal processing.

### This experiment differed from other context dependent experiments

Consistent with behavior, we found no support in the activity of hippocampal ensembles for a rule-specific code. These data are in stark contrast to previous experiments that presented animals with distinct behavioral contexts (e.g. Markus et al., 1995; McKenzie et al., 2014). However, it is important to note the difference between the contextual cues in this experiment and those in previous experiments. Those experiments that did observe segmentation used spatially distinguishable contexts (Komorowski et al., 2013b, 2009; McKenzie et al., 2014) or overt external cues to discriminate the context, such as an ambient sound, odor, or object (McNaughton et al., 1996; Zaremba et al., 2017). The task in the present experiment was specifically designed to be devoid of such ambient cues, as a neural response to that cue could be mistaken for a new context signal, providing false evidence for event segmentation. Other tasks generated contexts consisting of separate behavioral tasks or event sequences in the same physical space (Bower, Euston, & McNaughton, 2005; Ferbinteanu & Shapiro, 2003; Markus et al., 1995). But in those paradigms context was designated by overt changes in the physical behavioral sequences, and reward locations differed between contexts (Kobayashi et al., 1997; Markus et al., 1995). Thus, trajectory dependent firing ‘splitting,’ or cue responsivity may have been responsible for the context signal in these experiments (Ferbinteanu & Shapiro, 2003; Grieves, Wood, & Dudchenko, 2016). As the spatial layout of this task was held constant across the two rule conditions, there were no systematic changes in the animal’s trajectory, which could explain the absence of an alternating neural code.

A previous study found that in rats successfully performing alternation behaviors, alternation in the hippocampal code was not always apparent, and was sensitive the location of the reward itself (Bower et al., 2005). This study suggests that the shaping procedure in rodent experiments has a profound effect on context disambiguation in the hippocampus (Bower et al., 2005). Therefore, we hypothesize that rats may benefit from the opportunity to first explicitly learn two alternating rule conditions through training with an explicit contextual cue, such as wall color, or ambient sound. Rats might then continue to track the alternating context even after the ambient contextual cues are removed. This might also reveal a hippocampal representation of the two contexts and provide evidence for event segmentation when only boundary cues segment context. On the other hand, the hippocampus may code temporal context in a unique manner, and that other regions such as the Lateral Entorhinal Cortex may have shown segmentation in this task (Tsao et al., 2018a).

### These data add to a growing literature that describes temporal drift

Hippocampal populations showed robust drift through time both across seconds as well as minutes. These data add to a growing body of literature that suggests that hippocampal representations change across, and are sensitive to time. Importantly, this type of temporal drift has been observed in experiments employing a variety of recording methodologies (Cai et al., 2016; Mankin et al., 2015; Manns et al., 2007; Mau et al., 2018; Rubin, Geva, Sheintuch, & Ziv, 2015; Ziv et al., 2013). Hippocampal drift has also been observed in both hippocampal dependent tasks as well as tasks with no mnemonic demand (E. A. Mankin et al., 2012; Manns et al., 2007; see also: Tsao et al., 2018b). This experiment is to our knowledge the first direct evidence that population drift is uncorrelated with waveform drift in chronic tetrode recording preparations. Further suggesting that these time signals are unlikely to be a recording artifact, many units observed in the Lateral Entorhinal Cortex by Tsao et al. showed slow changes in spike rate that repeated multiple times and were triggered by entry into a new environment (Tsao et al., 2018b). Because drift is observed with recording techniques ranging from tetrode recording to calcium imaging to immediate early gene expression, this reduces the possibility that it is simply a measurement artifact - all of the recording methodologies would have to have independent artifacts that happen to produce the same results. This raises the possibility that population drift reflects a functional correlate of hippocampal processing that occurs continually even under no mnemonic demand, or when behavior is clamped.

A drifting representation of context supports the behavioral results in this experiment, and suggest that hippocampal processing contributed to the flexible behavior that was observed. This representation was supported on the individual unit level through firing fields centered on sampling events that were specific to position and object, but that peaked around a window of trials over the experimental session. Thus, drift occurred both in neurons coding for places and objects as well as in units without obvious place or object selectivity. Crucially, while some of these firing fields remained stable, across the population there was a spectrum of drift rates (Figure 6). The drifting representation of context is useful, as it provides a continuous dimension for relating experiences and is a likely mechanism for tracking the temporal relationships of events across many scales of time (Cai et al., 2016; Howard Eichenbaum, 2017). This may have enabled rats to forget past rule conditions and learn new ones by providing a mechanism for separating old experiences from more recent experiences. Specifically, this drift would enable rats to both associate very recent trials occurring in the same block and disassociate distant trials occurring in the previous block. Drift in the hippocampus may therefore enable the rapid learning that occurred around the beginning of each block.

### Was the hippocampus really insensitive to the event segmenting cues?

If the animals were generating new contextual representations at the onset of each block of trials, one might hypothesize that the boundary cue may have impacted the hippocampal code. Recent modeling suggests that event segmentation may occur when the actor detects shifts in the latent causes governing the set of rules in a given context, and that the hippocampus is necessary to properly assign a new context (Gershman, Monfils, Norman, & Niv, 2017; Gershman & Niv, 2010). Indeed the boundary cue, in this case the prolonged delay and removal from the task environment, did impact the rats’ expectations, as no rat showed response perseveration on the first trial of a block. fMRI studies in humans suggest this effect is caused by increased hippocampal activation coinciding with recognition of an event boundary (Baldassano et al., 2017; Swallow et al., 2011). This boundary effect has been recently replicated in the rodent hippocampus during two paradigms where space served to contextualize experience (Bulkin et al., 2018; Place et al., 2016). In this experiment, there was an increase in drift between trials separated by a block boundary, but the increased separation in representations was completely accounted for by the passage of time between blocks. This result suggests that removal from the environment had no impact on the hippocampal representation. However an alternative explanation is that time was sufficient to separate the hippocampal representation of context and promote a change in the rats’ expectations. Perhaps the temporal delay between blocks somehow inhibited the identification of a discrete change in the latent cause governing the reward contingency. Under this interpretation other cues may separate contexts as well if they were only given the right opportunity. Future experiments could dissociate the impact of each of these cues by imposing either the prolonged delay or the removal from the environment alone, or both in combination to observe different levels of separation in the hippocampal representation across time.

### A combined model of event segmentation and drift

In this study we examined the hippocampus for a neurophysiological signature of event segmentation in a behavioral task with a clear event boundary (removal from the testing environment). The assumption in designing the experiment was that it would be advantageous to map behaviorally-similar blocks of trials onto distinct cognitive maps and that the neural code would alternate across blocks of trials. As discussed above, there might be some methodological change to the experiment that would have enabled a successful event segmentation strategy. But another reason we failed to observe neural evidence for event segmentation is that our a priori expectations about how event segmentation would manifest at the resolution of individual neurons in the hippocampus may have been incorrect.

In humans, there is an active literature on how people segment continuous experiences, such as movies of real world scenes or radio plays (Zacks, Kurby, Eisenberg, & Haroutunian, 2011; Lositsky et al., 2016; Baldassano et al., 2017). Participants place event boundaries where they note meaningful changes in the ongoing structure of the world. One of the key findings from this literature is that people can segment boundaries at a range of time scales (Kurby & Zacks, 2008, 2011). For instance, suppose participants identify a segment several minutes long in a scene where a person makes a salad. At the same time people can also identify shorter segments within the larger segment corresponding to steps such as retrieving ingredients from the refrigerator, washing the vegetables, or chopping the vegetables. Indeed different environmental inputs in the real world change at many different rates, reflecting the complex multiscale structure of our world (Sreekumar, Dennis, Doxas, Zhuang, & Belkin, 2014; Voss & Clarke, 1975). Furthermore, fMRI evidence suggests different cortical regions may segment continuous experience into event segments at varying timescales (Baldassano et al., 2017). How might the hippocampus support the formation of event segments and the organization of segments across time?

One possibility is that hippocampal time cells possess all the properties necessary to support behavioral event segmentation. Critically, time cells respond at a wide range of delays tiling intervals that have been studied thus far up to at least tens of seconds (Salz et al., 2016). Furthermore, time cell sequences may be specific to certain past events, such as presentation of a specific odor (MacDonald, Carrow, Place, & Eichenbaum, 2013; Terada, Sakurai, Nakahara, & Fujisawa, 2017). Time cells may also track time at multiple time scales from minutes, to hours, to even days (Mau et al., 2018). These properties together suggest time cells may support our ability to bridge the gap between events at multiple scales to relate those experiences. A time cell sequence could provide a common temporal context that spans any boundary that occurred between experiences, thus providing a mechanism to relate those two experiences. Conversely, activity of overlapping time cells would also provide specific details about before and after a boundary has occurred, providing a mechanism to contrast those experiences.

As has been previously proposed, the hippocampus may provide pointers that index past events, rather than storing the content of those events per se (Teyler & DiScenna, 1986). At the ensemble level, pointers that drift across time enable the pointer system to reflect the temporal organization of events across time. In this way, temporally modulated drift coupled with the hippocampal place code enables a spatiotemporal scaffold onto which memory for events are stored.

## Acknowledgements

This project was originally conceived and designed in part by Dr. Howard Eichenbaum. Sadly, he passed away before the data were fully analyzed and therefore was unable to evaluate this manuscript in final form. The authors gratefully acknowledge his many contributions to this experiment, as well as the inspiration we’ve derived from his leadership. We would also like to thank Dr. Chris Keene for performing the initial pilot work for this experiment, and Dr. Michael Hasselmo for the close support and engaging discussions concerning this manuscript.

